# Precisely control mitochondria with light to manipulate cell fate decision

**DOI:** 10.1101/469668

**Authors:** Patrick Ernst, Ningning Xu, Jing Qu, Herbert Chen, Matthew S. Goldberg, Victor Darley-Usmar, Jianyi J. Zhang, Brian O’Rourke, Xiaoguang Liu, Lufang Zhou

## Abstract

Mitochondrial dysfunction has been implicated in many pathological conditions and diseases. The normal functioning of mitochondria relies on maintaining the inner mitochondrial membrane (IMM) potential (*a.k.a.* ΔΨ_m_) that is essential for ATP synthesis, Ca^2+^ homeostasis, redox balance and regulation of other key signaling pathways such as mitophagy and apoptosis. However, the detailed mechanisms by which ΔΨ_m_ regulates cellular function remain incompletely understood, partially due to difficulty of manipulating ΔΨ_m_ with spatiotemporal resolution, reversibility, or cell type specificity. To address this need, we have developed a next-generation optogenetic-based technique for controllable mitochondrial depolarization with light. We demonstrate successful targeting of the heterologous Channelrhodopsin-2 (ChR2) fusion protein to the IMM and formation of functional cationic channels capable of light-induced selective ΔΨ_m_ depolarization and mitochondrial autophagy. Importantly, we for the first time show that optogenetic-mediated mitochondrial depolarization can be well-controlled to differentially influence the fate of cells expressing mitochondrial ChR2: while sustained moderate light illumination induces substantial apoptotic cell death, transient mild light illumination elicits cytoprotection *via* mitochondrial preconditioning. Finally, we show that Parkin overexpression exacerbates, instead of ameliorating, mitochondrial depolarization-mediated cell death in HeLa cells. In summary, we provide evidence that the described mitochondrial-targeted optogenetics may have a broad application for studying the role of mitochondria in regulating cell function and fate decision.

## INTRODUCTION

Mitochondria lie at the crossroad of cellular metabolic and signaling pathways and thus play a pivotal role in regulating many intracellular processes such as energy production, reactive oxygen species (ROS) production, Ca^2+^ handling, and cell fate decision (1). Not surprisingly, defects of mitochondrial function have been implicated in a variety of human maladies such as aging, cardiovascular disease, cancer, and neurodegeneration (2). The normal functioning of mitochondria largely relies on maintaining the inner membrane potential (ΔΨ_m_) to preserve the protonmotive force necessary to drive oxidative phosphorylation and redox balance. Interestingly, while profound depolarization of ΔΨ_m_ is detrimental and induces cell injury (3), partial mitochondrial dissipation can be cytoprotective (4, 5). The detailed mechanism underlying the differential regulatory role of ΔΨ_m_ on cell function and fate remains incompletely understood, partially due to the difficulty of controlling ΔΨ_m_ or mitochondrial membrane permeability (MMP) with spatiotemporal resolution, reversibility, and with cell type specificity in an experimental setting. Currently, manipulation of ΔΨ_m_ is mainly *via* pharmacological intervention, i.e. using chemicals to uncouple the mitochondria (4, 5) or to induce permeability transition pore (mPTP) opening (6). However, pharmacological approaches often cause unknown side effects and lack the ability to probe spatiotemporal domains. Several research groups, including us, use laser flashes to trigger local or cell wide mitochondrial depolarization *via* the photooxidation mechanism (7–10). A limitation of this approach is that it requires high-energy laser illumination, which may induce uncontrollable oxidative stress and unpredictable outcomes. A new approach capable of precisely controlling MMP or ΔΨ_m_ is highly desired for advancing mitochondrial research.

Optogenetics has recently emerged as a technique that utilizes genetically-encoded light-sensitive ion channels, such as channelrhodopsin 2 (ChR2) (11–14) or halorhodopsin (NpHR) (15, 16), to precisely and remotely manipulate the activity of cells or animals (11, 13). ChR2 is a seven-transmembrane domain protein that contains the light-isomerizable chromophore all-trans-retinal (17). When excited by blue light, the *all-trans* retinal undergoes *trans-cis* isomerization (18), activating non-selective cationic channels and inducing depolarizing inward currents. Due to its high spatiotemporal resolution, ChR2-based optogenetic techniques have evolved as a powerful tool in basic and translational research. For example, ChR2 has been expressed in neurons (19) to monitor and control their membrane potential, intracellular acidity, Ca^2+^ influx, and other cellular processes (see ref (14) for review). In addition, ChR2 has been used to manipulate the membrane excitability of cardiomyocytes (20–22), skeletal muscle cells (23) and cell lines expressing voltage-gated ion channels (24, 25).

In addition to remote control of excitable cells, optogenetic techniques have been used to induce and study cell death. For instance, Yang *et al.* showed that an optogenetic approach can induce apoptotic cell death in human glioma cells *via* depolarization of cell membrane potential and influx of Ca^2+^ (26). In another study, Hill *et al.* reported a two-photon chemical apoptotic targeted ablation technique (27). By combining this technique with organelle-targeted fluorescent proteins and biosensors, the authors successfully achieved precise ablation of specific populations of neurons in the mouse brain. Of note, none of these published optogenetic approaches directly target the inner mitochondrial membrane (IMM) permeability to induce cell death. We hypothesize that functional expression of ChR2 in cellular IMM forms cationic channels capable of light-induced controlled ΔΨ_m_ depolarization and manipulation of cell function and fate. We test this hypothesis in a variety of cell types and our results show that the mitochondrial leading sequence (MLS) of ABCB10, an ATP binding cassette transporter, effectively targeted ChR2 into IMM. We demonstrate that this next-generation optogenetic approach can induce controlled ΔΨ_m_ depolarization that influences cell functions such as apoptosis and mitophagy at high spatial resolution.

## MATERIALS AND METHODS

### Cell culture

HeLa and H9C2 cells were obtained from ATCC and cultured in Dulbecco’s Modified Eagle Medium (Gibco) supplemented with 10% (v/v) Fetal Bovine Serum (Gibco) and 2 mM L-Glutamine (Gibco). Cells were maintained in 5% CO_2_ at 37 °C with confluence between 10% and 80%. hiPSC-CMs were reprogrammed from human cardiac fibroblasts as described previously (28) and maintained on Matrigel Membrane Matrix (Thermo Fisher Scientific) in mTeSR™ medium (Stem Cell Technologies) until 75% confluency. CM differentiation was performed by using a small molecule–based protocol as described previously (29, 30). Beating hiPSC-CM began to appear 9-12 days after differentiation was initiated.

### Plasmid construction

The ROMK and ABCB10(1-140AA) (named ABCB for simplification) MLS were synthesized by GeneArt (Life Technologies). pCMV-myc-mito-GFP vector was purchased from Life Technologies (catalog number V82220), and pLenti-EF1a-hChR2(H134R)-EYFP-WPRE was a gift from Karl Deisseroth (Addgene plasmid # 20942). To construct the CMV-myc-mito based plasmids (e.g. pCMV-mito-ChR2-eYFP), pCMV-myc-mito-GFP vector was digested with SalI and XbaI and ligated with ChR2-eYFP fragment obtained from pLenti-EF1a-hChR2(H134R)-EYFP-WPRE vector *via* PCR. The ROMK based plasmid was constructed by replacing the mito sequence with the ROMMK sequence. The pCAG-ABCB-ChR2-eYFP and pCAG-ABCB-eYFP vectors were constructed using the Gibson Ligation Kit following the manufacturer’s instructions. The sequences of constructed vectors were confirmed by DNA sequencing at the University of Alabama at Birmingham Heflin Center Genetics Core.

### Adenovirus production

The gene of interest (i.e. CMV-ABCB140-ChR2-YFP-ER) was PCR amplified from vector pCAG-CMV-ABCB-ChR2-eYFP using blunt-end primers. This gene fragment was cloned into the pENTR-D-TOPO vector purchased from Life Technologies (catalog number K2400) to generate the adenovirus entry vector. The entry vector sequence was confirmed using the M13 Forward and M13 Reverse primers. Adenoviral expression vector was constructed by LR recombination using the ViraPower™ Adenoviral Gateway™ Expression Kit (Life technologies catalog number K4930). To produce adenovirus, the expression vector was digested with PacI, purified, and then transfected into HEK 293A cells pre-seeded in T-flasks using Lipofectamine 2000. Cells were cultured until visible regions of cytopathic effect were observed in approximately 80% of the culture. Cells were then detached from the flask bottom and lysed *via* three successive freeze/thaw cycles to release the crude adenovirus. To amplify the adenovirus, 293A cells were infected with the crude adenovirus at a MOI of approximately 5. High titer adenovirus was harvested as described above when the 293A cells were floating or lightly attached to the culture dish, 2-3 days after infection.

### Confocal imaging and colocalization analysis

Plasmids containing ChR2-eFYP or eYFP with various MLS (e.g., mito, ROMK and ABCB) were transfected to H9C2 or HeLa cells cultured on a 15mm glass-bottom dish using the Lipofectoamine 3000 transfection reagent. hiPSC-CMs were transduced with the adenovirus containing ABCB-ChR2-eYFP at MOI of 50. 48 hours after transfection/transduction, cells were loaded with MitoTracker Deep Red (250 nM) for 30 minutes. The 515nm argon laser and 635nm LD laser lines of an Olympus FV1000 confocal microscope were used to image eYFP and MitoTracker, respectively. Images were processed offline using ImageJ software (Wayne Rasband, NIH) for colocalization analysis.

### Light illumination-induced targeted mitochondrial depolarization

The ABCB-ChR2-eYFP or ABCB-eYFP-expressing H9C2 cells were loaded with mitochondrial membrane potential fluorescent dye TMRM (20 nM) and illuminated with 475nm LED pulses (5 mW/mm^2^). The blue light was delivered by a 10X objective lens coupled to a LED driver, which was controlled by a Myopacer stimulator (IonOptix, Westwood, MA). The fluorescence of TMRM during light illumination was recorded using an Olympus FV1000 confocal microscope and analyzed using ImageJ software.

### Cell death and cytotoxicity assay

Cells were illuminated by LED light of different duration and irradiance (as indicated in the corresponding figure legends). Right after light treatment cells were detached with 0.25% Trypsin-EDTA, stained with Trypan Blue, and counted using a hemocytometer as described previously (25). The cytotoxicity and cell viability were also measured using the LDH assay and AlamarBlue (AB) assay, respectively. For the LDH assay, at the end of light illumination, 200 μl cell culture medium was removed and added to a 96 well plate, 100 μl of the reaction mixture was added to each well and the plate was incubated for 30 minutes at room temperature, protected from light. The absorbance was measured at 490nm and 680nm using a microplate reader. LDH activity was calculated following the manufacture’s instruction. For the AB assay, 10 μl AlamarBlue reagent was directly added to cells in 100 μl culture medium. Cell cultures were incubated for 4 hours in an incubator (37 °C, protected from light). The absorbance was measured at 570nm using a spectrometer.

### Immunocytochemistry

Cells were fixed in 4% Formaldehyde in PBS and then treated with PBS containing 10% goat serum and 0.3% Triton® X-100 to block non-specific staining. Cells were then stained with the following primary antibodies: ChR2, TOMM20, LC3, or Cytochrome C at a ratio of 1:200. After overnight room temperature incubation, secondary antibodies AlexaFluor 555 and/or AlexaFluor 647 were diluted in 1% BSA, 1% goat serum, 0.3% Triton® X-100 in PBS at a ratio of 1:200, and then added to the cells. Coverslips were mounted on slides and imaged using an Olympus FV1000 confocal microscope.

### Parkin translocation assay

HeLa cells were co-transfected with plasmid encoding mCherry-tagged Parkin (31) (Addgene plasmid #23956;) and encoding either ABCB-ChR2-YFP or ABCB-YFP using the Neon Transfection System (Invitrogen) and seeded in 35 mm glass-bottom dishes (MatTek, Ashland, MA). Two days after transfection cells were subjected to LED illumination (0.5 mW/mm^2^, 24 hours). Confocal imaging of YFP and mCherry expression and localization was done immediately before treatment, and after 4, 8, and 24 hours of treatment.

### Detection of the colocalization of mitochondria and lysosomes

HeLa cells were transduced with ABCB-ChR2-eYFP adenovirus and transfected with mCherry-Parkin plasmid DNA. Forty-eight hours later LED stimulation (0.5 mW/mm^2^) of the cells began. After 24 hours of stimulation, cells were stained with 50 nM Lysotracker Deep Red at 37 °C for 30 minutes, followed by three washes. Fluorescent images were observed with confocal microscopy. Green fluorescence (i.e. YFP) represents mitochondria and red (LysoTracker) represents lysosomes. The colocalization of mitochondria and lysosomes indirectly indicates mitophagy. All light stimulation was done at a frequency of 4 Hz with 90 ms pulses. Un-illuminated mock-transfected and ABCB-ChR2-eYFP expressing cell cultures were used as control.

### Mitochondrial preconditioning

HeLa cells were transduced with ABCB-ChR2-eYFP adenovirus and transfected with mCherry-Parkin plasmid DNA. Forty-eight hours later cultures were subjected to their respective treatments: no light stimulation for control group, 2 hours of no light stimulation followed by 6 hours of stimulation at 4 mW/mm^2^ for stimulation-only group, and 2 hours of stimulation at 0.2 mW/mm^2^ followed by 6 hours of stimulation at 4 mW/mm^2^ for pre-conditioning group. Cell cultures were then stained with Mitoview 633 (25 nM) at 37 °C for 15 minutes, washed of dye, and then immediately followed by confocal imaging. All light stimulation was done at a frequency of 4 Hz with 90 ms pulses.

### Flow cytometry

Cells were trypsinized and washed twice with PBS before resuspension in staining buffer (10 mM HEPES/NaOH (pH 7.4), 140 mM NaCl, 2.5 mM CaCl_2_) containing MitoView 633 (25 nM) and stained following the manufacture’s instruction. Cells were then washed twice with PBS + 1% BSA before analysis with a BD LSR II Flow Cytometer (BD Biosciences, San Jose, CA).

### Materials

Mouse monoclonal antibodies for GFP, ATP5A, TOM20 and LC3 were purchased from Abcam (Cambridge, MA). Mouse monoclonal ChR2 antibody was obtained from Progen (Heidelberg, Germany). Oregon Green 488 anti-rabbit IgG, MitoSox, TMRM, and MitoTracker Deep Red FM were purchased from Life Technologies (Carlsbad, CA). MitoView 633 was purchased from Biotium (Fremont, CA). All other reagents were obtained from Sigma (St Louis, MO).

### Statistics

Analysis and presentation of the data were performed using OriginLab software (Northampton, MA). Comparisons between groups were performed using paired or unpaired 2-tailed Student’s t test. Data were considered significantly different at *p* < 0.05 or below. Results are presented as mean ± SEM.

## RESULTS

### Targeting ChR2 to mitochondria

The MLS of cytochrome *c* oxidase subunit VIII (*a.k.a.* mito) has been widely used to import proteins such as GFP (32) or pericam (33) to mitochondria. Therefore, we first examined its capability to target ChR2 to mitochondria. A mutant of ChR2, ChR2(H134R) (34), was fused with eYFP and inserted into the vector pCMV-myc-mito to generate plasmid mito-ChR2(H134R)-eYFP. The construct and the control vectors (mito-eYFP) were transfected to H9C2 cells, respectively. The expressions of ChR2-eYFP and eYFP in mitochondria were examined by staining mitochondria with Mitoracker® Deep Red and analyzing the colocalization of the green and red channels using confocal microscopy. Results show that eYFP was perfectly colocalized with mitochondria (Figure S1A) but ChR2-eYFP was not (Figure S1B). In addition, ChR2-eYFP appeared to be improperly processed by the endoplasmic reticulum (ER), implying that the size of the ChR2(H134R)-eYFP precursor might be impairing import. Thus, we next duplicated and triplicated the copy number of the mito sequence and constructed plasmid (mito)_2_-ChR2(H134R)-eYFP and (mito)_3_-ChR2(H134R)-eYFP, respectively. However, none of these modifications improved the import of ChR2(H134R)-eYFP to mitochondria (Figures S1C and S1D). We also studied the MLS of ROMK, a candidate of mitochondrial K_ATP_ channel located in the IMM (35). Similarly, ROMK MLS successfully imported eYFP into mitochondria but not ChR2-eYFP (Figure S1E).

ATP binding cassette (ABC) transporters comprise a large family of membrane translocases (36) and are found in all species from microbe to human (37). Of them, ABCB10 is an IMM erythroid transporter with an unusually long MLS (140 amino acids) (38). Studies showed that ABCB10 MLS could lead various membrane proteins to mitochondria (39), while its depletion targeted ABCB10 to the ER (38). This MLS was fused with eYFP or ChR2(H134R)-eYFP, to generate ABCB-eYFP and ABCB-ChR2(H134R)-eYFP vectors, respectively, which were then transfected into H9C2 cells. To our great satisfaction, ABCB-eYFP was exclusively localized in mitochondria (Figure 1A). Excitingly, ABCB10 MLS also successfully translocated the ChR2-eYFP precursor into H9C2 mitochondria, as demonstrated by the colocalization of green (eYFP) and red (MitoTracker) channels (Figure 1B). Importantly, this method of targeting ChR2 to the IMM did not significantly affect mitochondrial membrane potential and reactive oxygen species (ROS) levels (Figure S2). The disposition of the ChR2-YFP fusion protein in H9C2 mitochondria was further examined by immunostaining. As shown in figure 1C, the ChR2 protein expression, detected by the ChR2 antibody, colocalized well with MitoTracker. The capability of the ABCB10 MLS to translocate ChR2-eYFP to mitochondria was also confirmed in other cell types such as HeLa cells (Figure 1D) and human induced pluripotent stem cell-derived cardiomyocytes (hiPSC-CMs) (Figure 1E). Altogether, our data demonstrate successful mitochondrial ChR2 import and expression.

**Figure 1.**
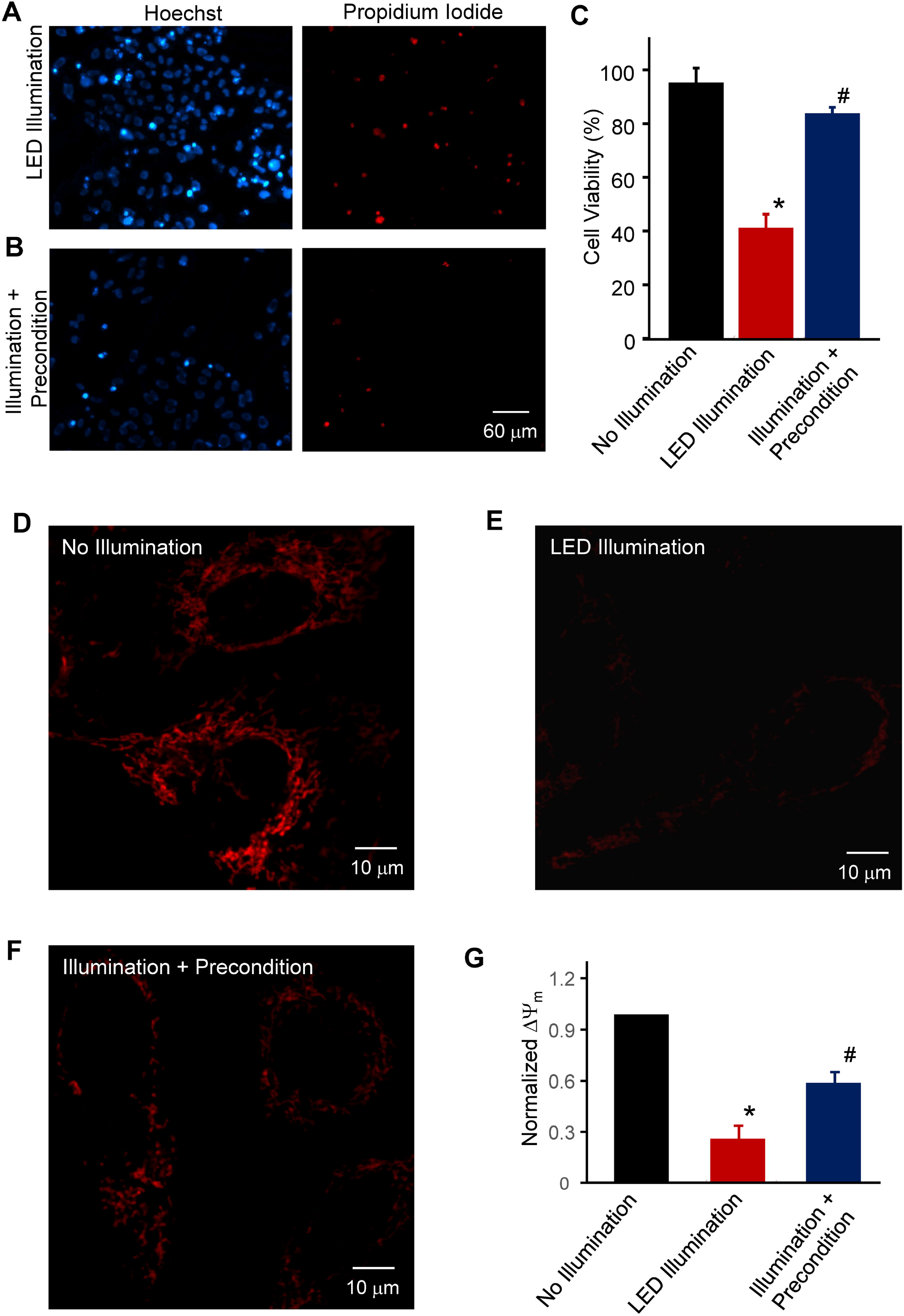
Representative confocal images showing intracellular localization of ChR2-eYFP or eYFP expression in different cell types. A): H9C2 cells expressing ABCB-eYFP; B): H9C2 cells expressing ABCB-ChR2-eYFP; C): Immunostaining of H9C2 cells expressing ABCB-ChR2-eFYP with ChR2 antibody; D): HeLa cells expressing ABCB-ChR2-eYFP; and E): hiPSC-CMs expressing ABCB-ChR2-eYFP. In all experiments cells were stained with MitoTracker Deep Red (250 nM). Yellow indicates colocalization of YFP (in A), ChR2-eYFP (in B, D, and E) or ChR2 (in C) and the mitochondrial marker.

### Light illumination-elicited targeted mitochondrial depolarization

After demonstrating mitochondrial ChR2 expression, we examined if light illumination could cause targeted mitochondrial depolarization in ABCB-ChR2-expressing cells. In the example shown in figure 2A, H9C2 cells expressing ABCB-ChR2 (fused with eYFP) were split into two zones with zone 1 receiving blue (475 nm) LED pulse illuminations and zone 2 as control (i.e. no LED illumination). The change of ΔΨ_m_ was measured with TMRM using confocal microscopy (Figure 2B), which showed that TMRM fluorescence faded significantly in the illuminated cells (i.e. zone 1) but did not change in the unilluminated cells (i.e. zone 2) (Figure 2C and Video S1). The averaged TMRM intensities before and after light illumination in the unilluminated and illuminated cells were summarized in figure 2D, showing that ΔΨ_m_ depolarized approximately 80% in the illuminated cells but remained polarized in the non-illuminated cells. The data suggest that mitochondrial optogenetics can induce mitochondrial depolarization with spatial resolution. In addition to impaired mitochondrial membrane potential, light illumination also caused increased ROS production in ABCB-ChR2 expressing cells, as compared with mock-transfected or ABCB-ChR2-expressing cells without blue light illumination (Figure S3). To verify that mitochondrial depolarization in the illuminated cells was caused by light-induced activation of IMM ChR2 channels instead of imaging laser-induced photooxidation and photobleaching, we performed two control experiments. We first examined the effect of the imaging laser alone on ΔΨ_m_ in ABCB-ChR2-expressing cells. As shown in figure S4A (and Video S2A), the confocal scanning laser itself had no effect on TMRM. However, turning on blue LED illumination led to significant ΔΨ_m_ depolarization (Figure S4B and Video S2B) similar to that shown in figure 2. In another experiment, we applied the same imaging and LED illumination as that used in the experiment shown in figure 2 to ABCB-eYFP-expressing cells. We found that neither eYFP nor TMRM fluorescent intensity changed upon LED illumination (Figure S5 and Video S3). Taken together, these data demonstrate functional expression of ChR2 in the IMM that allows light-induced spatially-selective mitochondrial depolarization.

**Figure 2.**
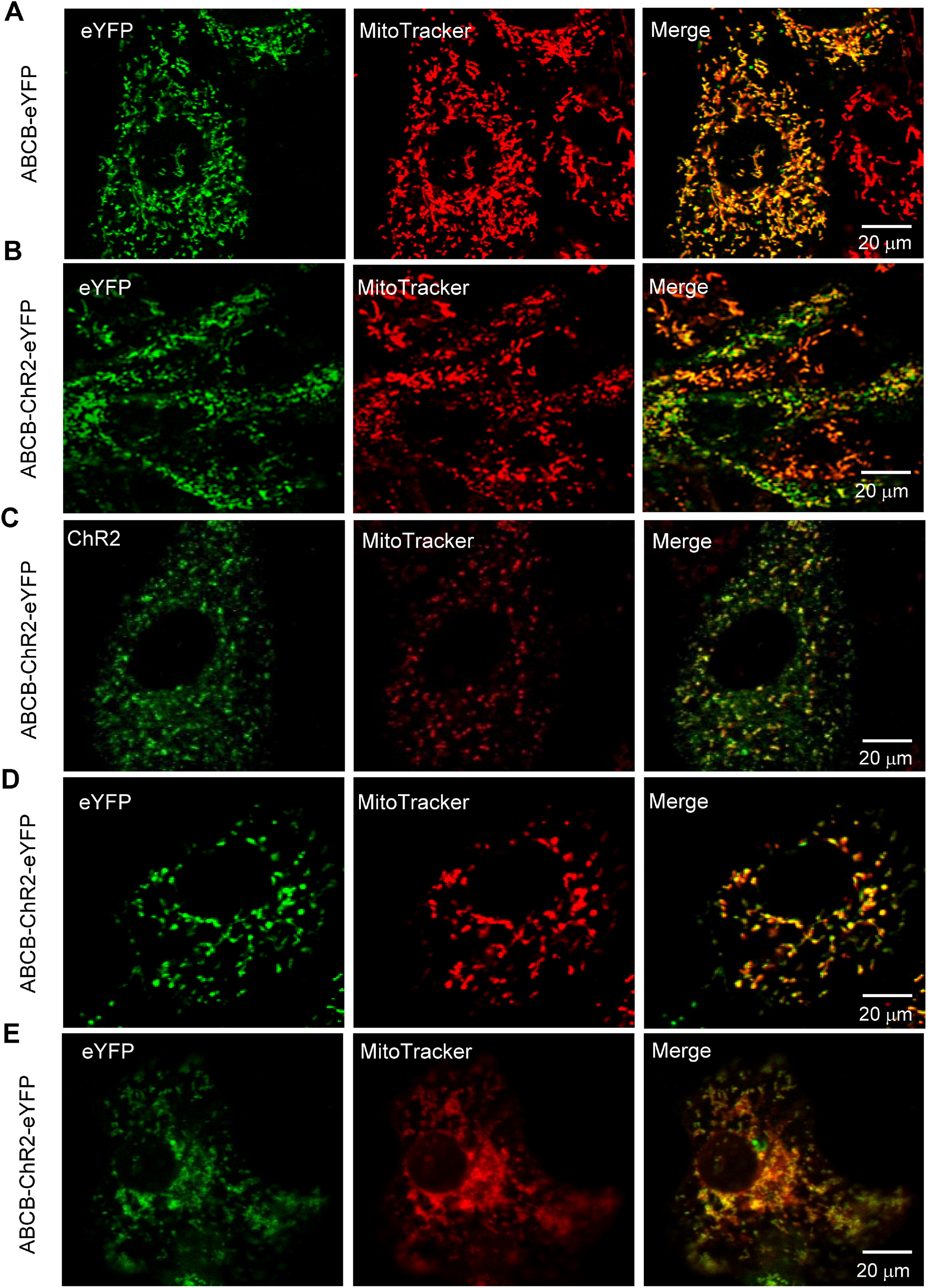
Mitochondrial-targeted optogenetics induces selective mitochondrial depolarization in H9C2 cells expressing ABCB-ChR2-eYFP. A): Schematic Figure Showing that cells in zone 1 were illuminated by LED (5 mW/mm^2^) and cells in zone 2 were not exposed to illumination; B) and C): Confocal images showing mitochondrial membrane potential (ΔΨ_m_), measured by TMRM fluorescent dye (20 nM), of cells in zone 1 and zone 2 before (B) and after (C) light illumination; D): Normalized ΔΨ_m_ in cells in the illuminated (i.e. zone 1) and un-illuminated (i.e. zone 2) zones before and after light illumination. 2-4 cells in each zone were analyzed in each cell culture and totally 4 cell cultures were examined.

### Mitochondrial optogenetic-mediated cell death and the underlying mechanisms

Next, we examined the effect of sustained light illumination on ABCB-ChR2-expressing cells. Twenty-four hours of moderate (0.5 mW/mm^2^) LED illumination had negligible effects on the viability of non-transfected and mock-transfected HeLa cells (Figure S6), as well as ABCB-eYFP-expressing HeLa cells, but caused significant cell death in ABCB-ChR2-expressing HeLa cells (Figure 3A), as measured by Trypan Blue staining. The LED illumination-induced cytotoxicity in ABCB-ChR2-expressing cells was confirmed by the lactate dehydrogenase (LDH) and AlamarBlue assays. As shown in figure 3B, cell viability dropped 80% in the light-illuminated ABCB-ChR2 cell cultures but only slightly in the ABCB-eYFP-expressing HeLa cells. Consistently, substantial cytotoxicity was detected in the light-illuminated ABCB-ChR2-expressing cells but not in the ABCB-eYFP-expressing cells (Figure 3C). Importantly, we found that pretreating cells with cyclosporine A (CsA, 1 μmol/L)), a selective mitochondrial permeability transition pore (mPTP) inhibitor, did not prevent the light-induced cell death (Figure 3D) and cytotoxicity (Figure 3E), indicating that the mitochondrial-targeted optogenetic-mediated cell death is independent of mPTP opening. This finding also denotes that ChR2 proteins formed mPTP-like ion channels in IMM. Of note, antioxidants such as MitoTEMPO and MitoQ failed to rescue cells from light-induced mitochondrial depolarization and cell death (Figure S7), suggesting that mitochondrial-derived ROS production and the ROS-induced ROS-release is not a crucial factor in mitochondrial optogenetic-mediated cytotoxicity.

**Figure 3.**
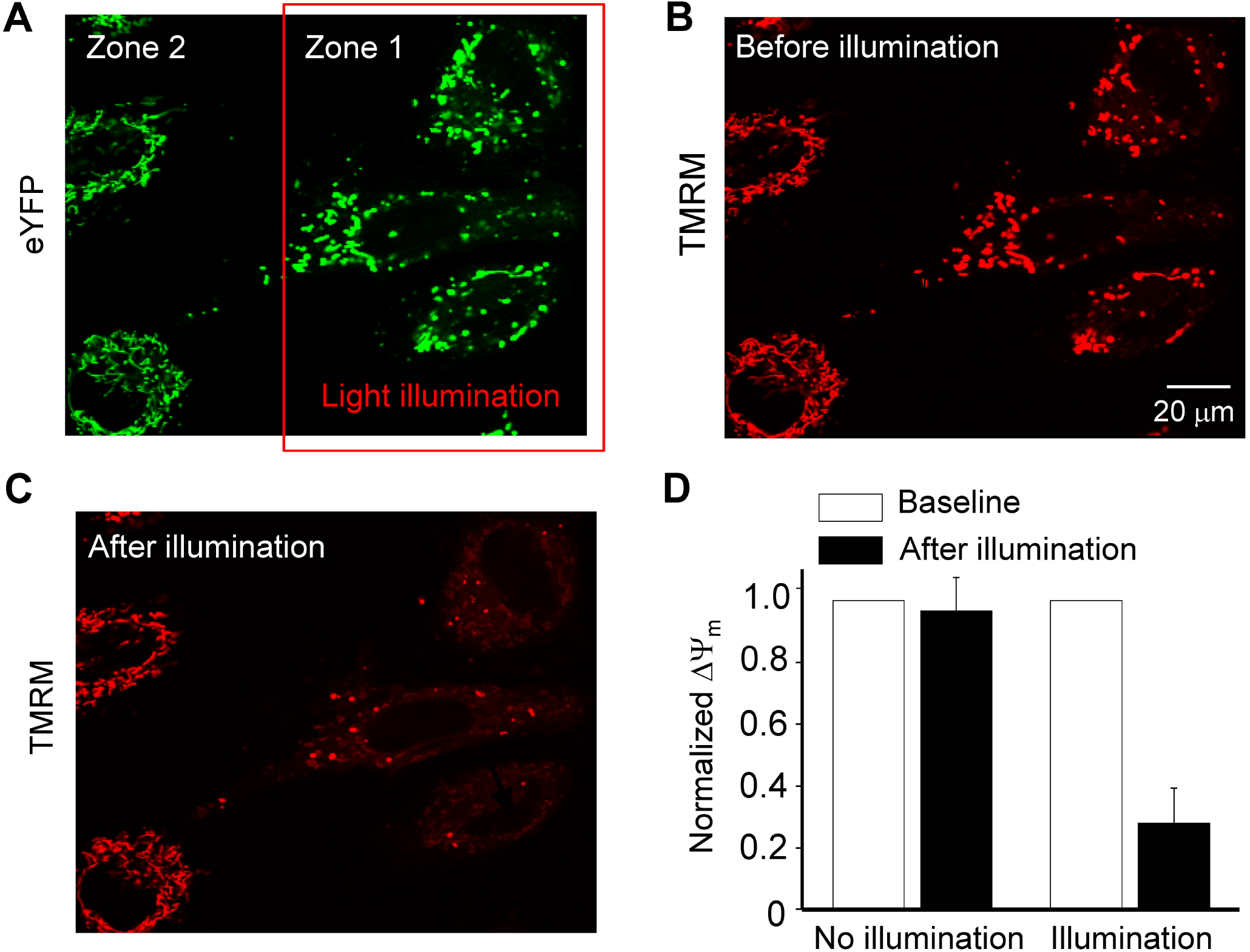
Mitochondrial-targeted optogenetics induces cell death in HeLa cells expressing ABCB-ChR2-eYFP. A): Representative transmitted light images of ABCB-ChR2-eYFP or ABCB-eYFP-expressing cells with or without LED illumination (0.5 mW/mm^2^, 24 hours); B) and C): Comparison of cell viability measured by LDH assay (B) and cytotoxicity measured by alamarBlue assay (C) in ABCB-ChR2-eYFP or ABCB-eYFP cells with or without LED illumination; D) and E): Effect of a mPTP blocker, CsA, on cell viability (D) and cytotoxicity (E) in ABCB-ChR2-eYFP-expressing or mock-transfected cell cultures with or without LED illumination. n represents the number of cell cultures.

We also examined the time and light irradiance dependence of optogenetic-mediated cell death. As shown in figure 4A, cell viability dropped nearly linearly within 12 hours of light (0.5 mW/mm^2^) illumination and tapered off slightly thereafter. The same light illumination had little effect on the viability of mock-transfected cultures (Figure 4A), consistent with results in figure S6. Regarding the effect of light intensity, we found that six hours of moderate (e.g., 1 mW/mm^2^ and 3 mW/mm^2^) light illumination caused minimal cell death in ABCB-ChR2-expressing cell cultures, as measured by Hoechst and Propidium Iodide (PI) staining (Figure S8). Increasing light intensity to 5 mW/mm^2^ or 7 mW/mm^2^ caused a substantial (62% and 85%, respectively) decrease in cell viability (Figures 4B and 4C). Importantly, light illumination at similar intensities (e.g., 3 mW/mm^2^ and 5 mW/mm^2^) did not induce detectable cytotoxicity in the mock-transfected cultures (Figures 4B, 4C and S8). When light intensity was further increased (e.g. >7 mW/mm^2^), illumination (24 hours) begun to cause cell death in these cells (e.g. illumination at 8 mW/mm^2^ decreased viability to 76%).

**Figure 4.**
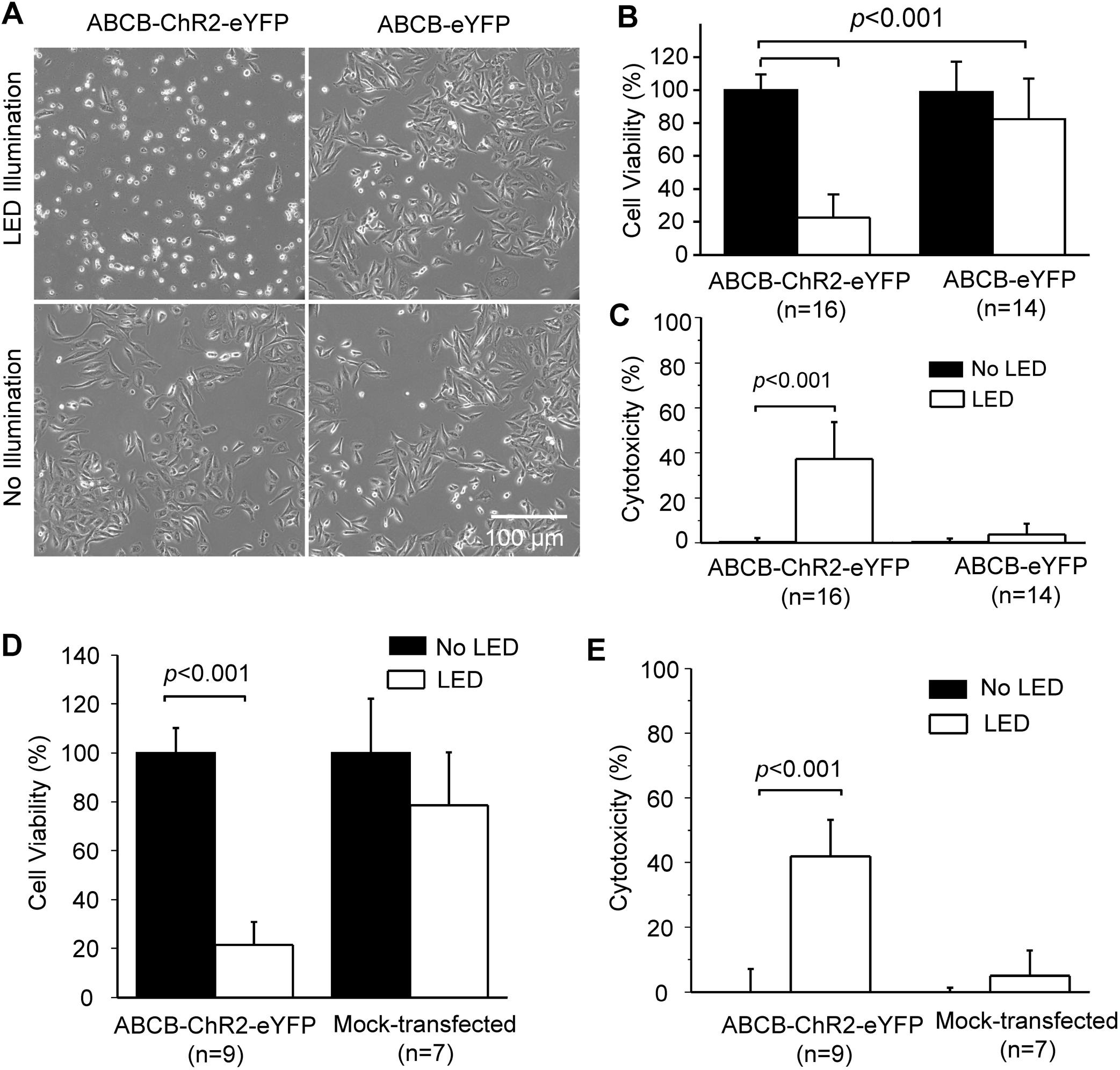
Time and light irradiance dependence of mitochondrial optogenetic-mediated cell death. A): Time dependence of the effect of moderate LED illumination (0.5 mW/mm^2^) on cell viability in the mock-transfected and ABCB-ChR2-eYFP-expressing HeLa cell cultures. Four cell cultures were repeated at each time point. Cell viability was determined using the trypan blue method; B): LED irradiance dependence of illumination-induced cell death. ABCB-ChR2-eYFP or mock-transfected HeLa cultures were illuminated by LED of various intensity for six hours. At least four cell cultures were repeated for each group. Cell death was measured by Propidium iodide (PI) staining; C): Representative images of PI staining in cultures illuminated by LED of different irradiance summarized in (B).

To explore the mechanistic pathway underlying the mitochondrial optogenetic-mediated cell death, we treated cells with an apoptosis inhibitor and a necroptosis inhibitor. Cell death analysis showed that Z-VAD-FMK (20μM), a pan-caspase inhibitor, substantially reduced light illumination-induced cell death, but 7-Cl-O-Nec-1 (100μM), a necroptosis inhibitor, had little effect on cell viability (Figure 5A), suggesting that the optogenetic-mediated light-induced cell death is likely through the caspase-dependent apoptotic pathway. We also noticed that Z-VAD-FMK-treatment significantly reduced ΔΨ_m_ depolarization in cell cultures exposure to light illumination (Figures 5B and 5C). The similar effect of caspase inhibition on mitochondrial depolarization in apoptotic cells has been reported in other studies (40, 41). The light illumination-elicited activation of the intrinsic apoptosis pathway in ABCB-ChR2-expressing cells was confirmed by cytochrome C release experiments. As shown in figure 5C, immunocytochemistry revealed that cytochrome C was colocalized with TOM20, a mitochondrial outer membrane protein, in both the mock-transfected cells and ABCB-ChR2-expressing cells not illuminated by LED light. However, light-illuminated ABCB-ChR2-expressing cells showed a punctate/diffuse cytochrome C staining, which was disparate from TOM20 immunostaining, indicating the release of cytochrome C into the cytosol (Figure 5D).

**Figure 5.**
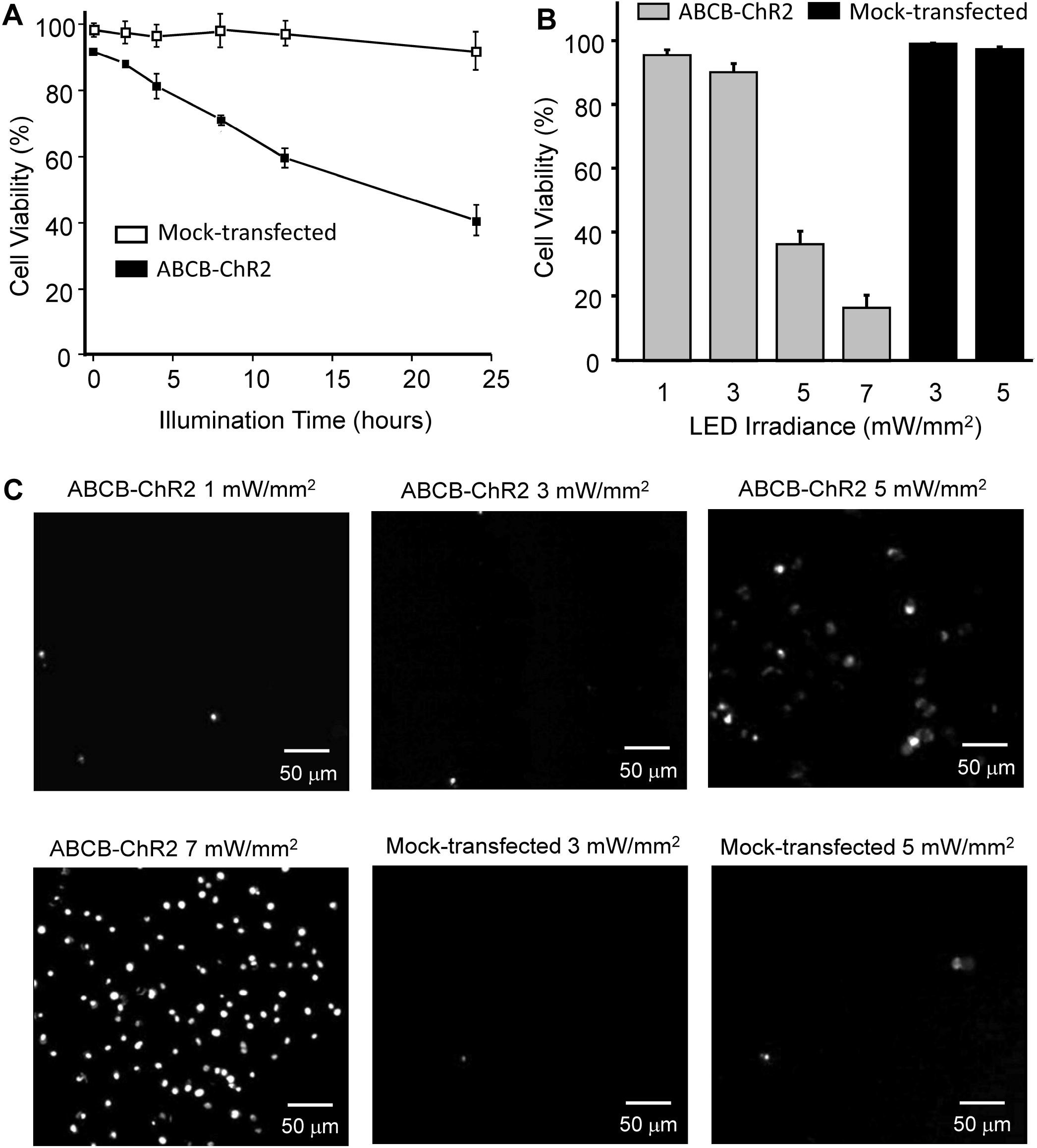
Mitochondrial optogenetics induces cell death *via* the apoptotic pathway. A): Z-VAD-FMK, a pan caspase inhibitor, substantially ameliorated light-elicited cell death in HeLa cells expressing ABCB-ChR2. 7-Cl-O-Nec-1, a necroptosis inhibitor, had no significant effect on the light-induced cell death. n represents the number of cell cultures; B): Representative flow cytometry analysis showing that Z-VAD-FMK alleviated light-induced mitochondrial depolarization in ABCB-ChR2-expressing cells; C: Summary of data showing the effect of Z-VAD-FMK on mitochondrial membrane potential (measured by MitoView) in cells exposed to light illumination; and D): Representative confocal images showing that light illumination (0.5 mW/mm^2^, 24 hours) induced cytochrome C release into the cytosol in ABCB-ChR2-expressing cells. *: P<0.05 *vs*. ABCB-ChR2 &No LED illumination. #: P<0.05 *vs*. ABCB-ChR2 &LED illumination. n=4 each group.

### Mitochondrial optogenetic-mediated mitochondrial autophagy

We next examined whether optogenetics-mediated mitochondrial depolarization and damage elicits mitochondrial autophagy (*a.k.a.* mitophagy). As HeLa cells do not express endogenous Parkin that is required for the canonical PINK1/Parkin mediated mitophagy pathway, we overexpressed mCherry-Parkin in ABCB-ChR2-eYFP-expressing HeLa cells, which were then subjected to LED illumination (24 hours, 0.5 mW/mm^2^). The cellular localization of Parkin was examined by confocal microscopy at different time points during light illumination. As shown in figure 6A, at the baseline (i.e. 0 hour) mitochondria (represented by eYFP fluorescence) are tubular like and mCherry-Parkin was clearly cytosolic and diffuse, as expected. After 4 hours of LED illumination, mCherry-Parkin began to translocate from the cytosol to some mitochondria, as indicated by colocalization of Parkin and eYFP. After 8 hours of LED illumination, mitochondria became fragmented and mCherry-Parkin translocated to most of the mitochondria. After 24 hours, most of the mCherry-Parkin was found to surround mitochondria, as shown in the enlarged view of the boxed region, demonstrating that Parkin was recruited to mitochondrial outer membranes. On the contrary, no evident mitochondrial translocation of Parkin was observed in the light-illuminated ABCB-eYFP-expressing cells (Figure S9A) and the un-illuminated ABCB-ChR2-eYFP-expressing cells (Figure S9B) after 24 hours. Furthermore, light-illuminated ABCB-ChR2-eYFP-expressing cells displayed less mitochondrial mass compared with other groups, especially after 24 hours, suggesting induction of Parkin-dependent mitophagy. The optogenetic-induced mitochondrial autophagy was confirmed by the aggregation of the autophagosome marker LC3 (Figure 6B, red) and the lysosome marker LysoTracker (Figure 6C, red) and their colocalization with mitochondria (represented by YFP). Neither marker aggregated and exhibited colocalization with YFP in the un-illuminated ABCB-ChR2-eYFP cells (Figures S9C and S9E) or the mock-transfected cells (Figure S9D). Together, these data demonstrate mitochondrial optogenetic-induced mitophagy.

**Figure 6.**
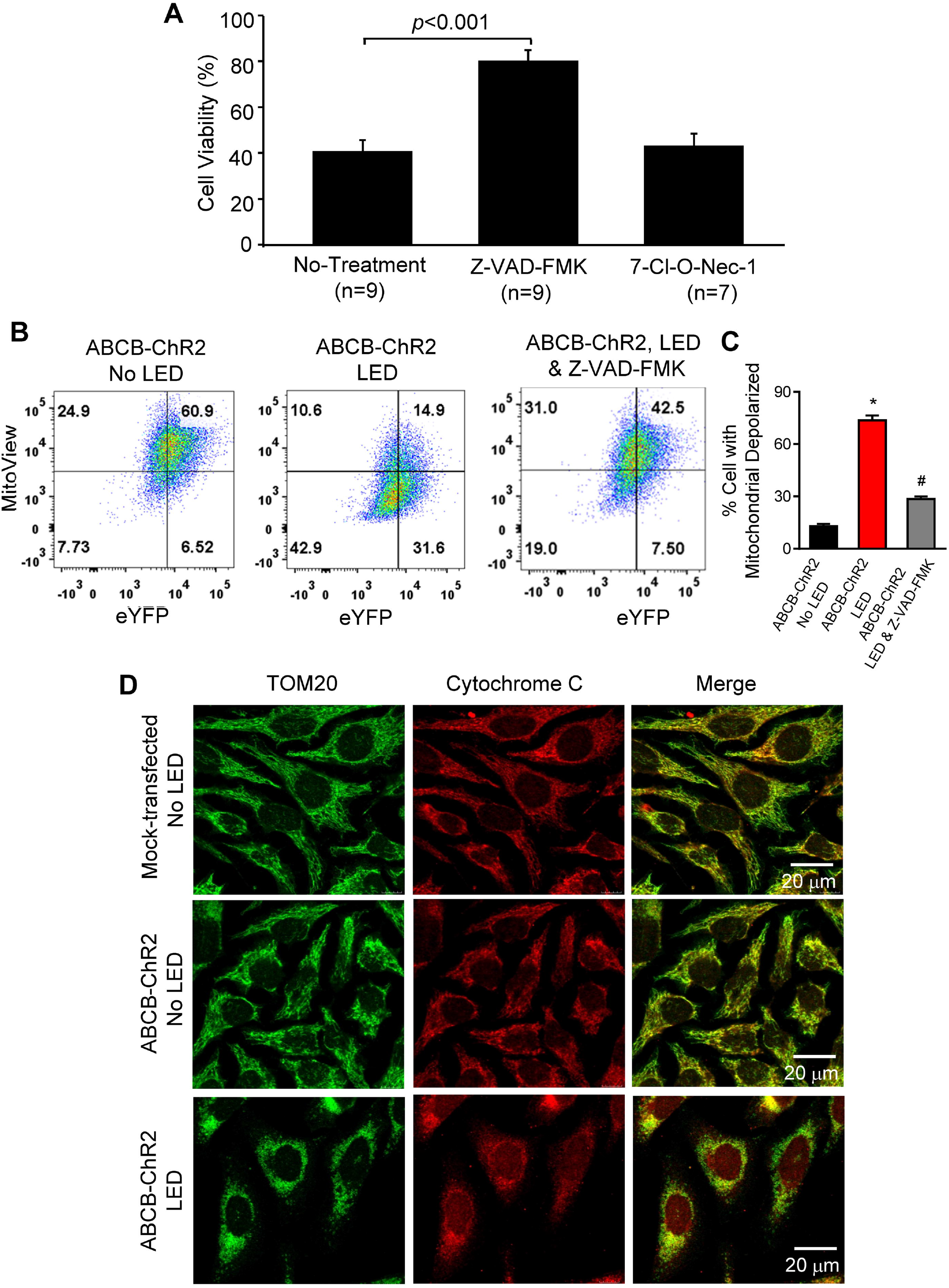
Light illumination induces mitochondrial autophagy in ABCB-ChR2-eYFP-expressing cells. A): Representative confocal imagines showing time-dependent localization of Parkin-mCherry in light-illuminated cells expressing ABCB-ChR2-eYFP and Parkin-mCherry. Images were taken at 0, 4, 8, and 24 hours during light illumination (0.5 mW/mm^2^). Red represents Parkin-mCherry, green represents eYFP, and overlay represents Parkin accumulation in mitochondria; B): Representative confocal imagines showing LC3 (red) aggregation and colocalization with mitochondria (represented by YFP) in illuminated (0.5 mW/mm^2^, 24 hours) ABCB-ChR2-eYFP expressing cells; and C): Representative confocal imagines showing lysosomes (red, represented by LysoTracker) aggregation and colocalization with mitochondria (represented by YFP) in illuminated (0.5 mW/mm^2^, 24 hours) ABCB-ChR2-eYFP expressing cells.

As activation of mitophagy is thought to be a cell survival mechanism by eliminating damaged mitochondria, we asked whether Parkin overexpression and activation of Parkin-mediated mitophagy pathway reduces optogenetic-mediated cell death. ABCB-ChR2-expressing HeLa cells, with or without Parkin expression, were illuminated with light (24 hours, 0.5 mW/mm^2^) and then subjected to ΔΨ_m_ and cell death analysis. As expected, expression of Parkin had little effect on ΔΨ_m_ (Figures 7A and 7B) in the absence of light illumination. However, Parkin overexpression surprisingly exacerbated light-induced mitochondrial depolarization (Figures 7A and 7B). In addition, although Parkin expression did not result in evident difference in cell viability in ABCB-ChR2-expressing cells immediately after 24 hours light illumination, significantly more cell death was observed in the Parkin-expressing HeLa cells 24 hours after stopping LED illumination (Figure 7C). The similar pro-apoptotic effect of Parkin activation has been reported in other studies (42, 43).

**Figure 7.**
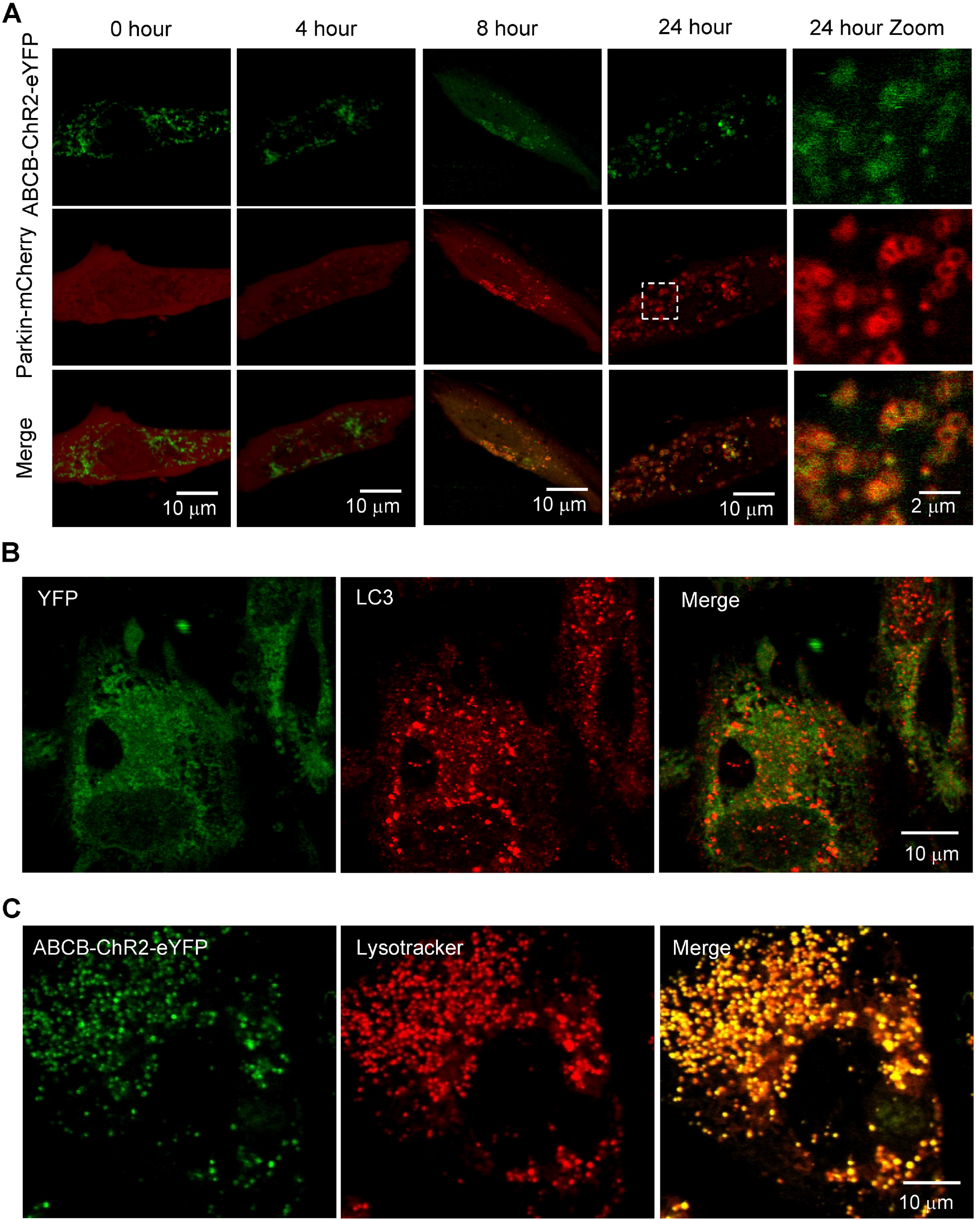
Parkin activation exacerbates mitochondrial optogenetic-mediated mitochondrial depolarization and cell death. A): Representative flow cytometry analysis showing mitochondrial membrane potential (measured by MitoView) in the illuminated (0.5 mW/mm^2^, 24 hours) or un-illuminated HeLa cells with or without Parkin overexpression; B): Summary of data showing that Parkin overexpression exacerbated light illumination-mediated mitochondrial depolarization in HeLa cells; and C): Parkin overexpression and activation did not influence HeLa cell viability immediately following light illumination (0.5 mW/mm^2^, 24 hours) but promoted cell death 24 hours later. 5 cell cultures were examined for each group. *: P<0.05 *vs*. No Parkin &No LED illumination. #: P<0.05 *vs*. No Parkin &LED illumination.

### Mitochondrial optogenetic-mediated preconditioning and cytoprotection

As previous studies indicated that mitochondrial preconditioning protects cells from stress, we examined if mitochondrial optogenetic-mediated partial mitochondrial depolarization reduces apoptotic cell death in ABCB-ChR2-expressing cells. As shown in figure 8A, 6 hours of moderate light (4 mW/mm^2^) illumination caused nearly 60% decrease in cell viability, which was consistent with data reported in figure 5. Intriguingly, cell viability in those cultures pretreated with 2 hours of mild (0.2 mW/mm^2^) light illumination was significantly higher compared with the non-preconditioned group (i.e. without pretreatment) (Figure 8B), as summarized in figure 8C. In addition, confocal imaging showed that light illumination caused 78% reduction in the mitochondrial membrane potential and preconditioning significantly alleviated mitochondrial depolarization (Figures 8D-8G).

**Figure 8.**
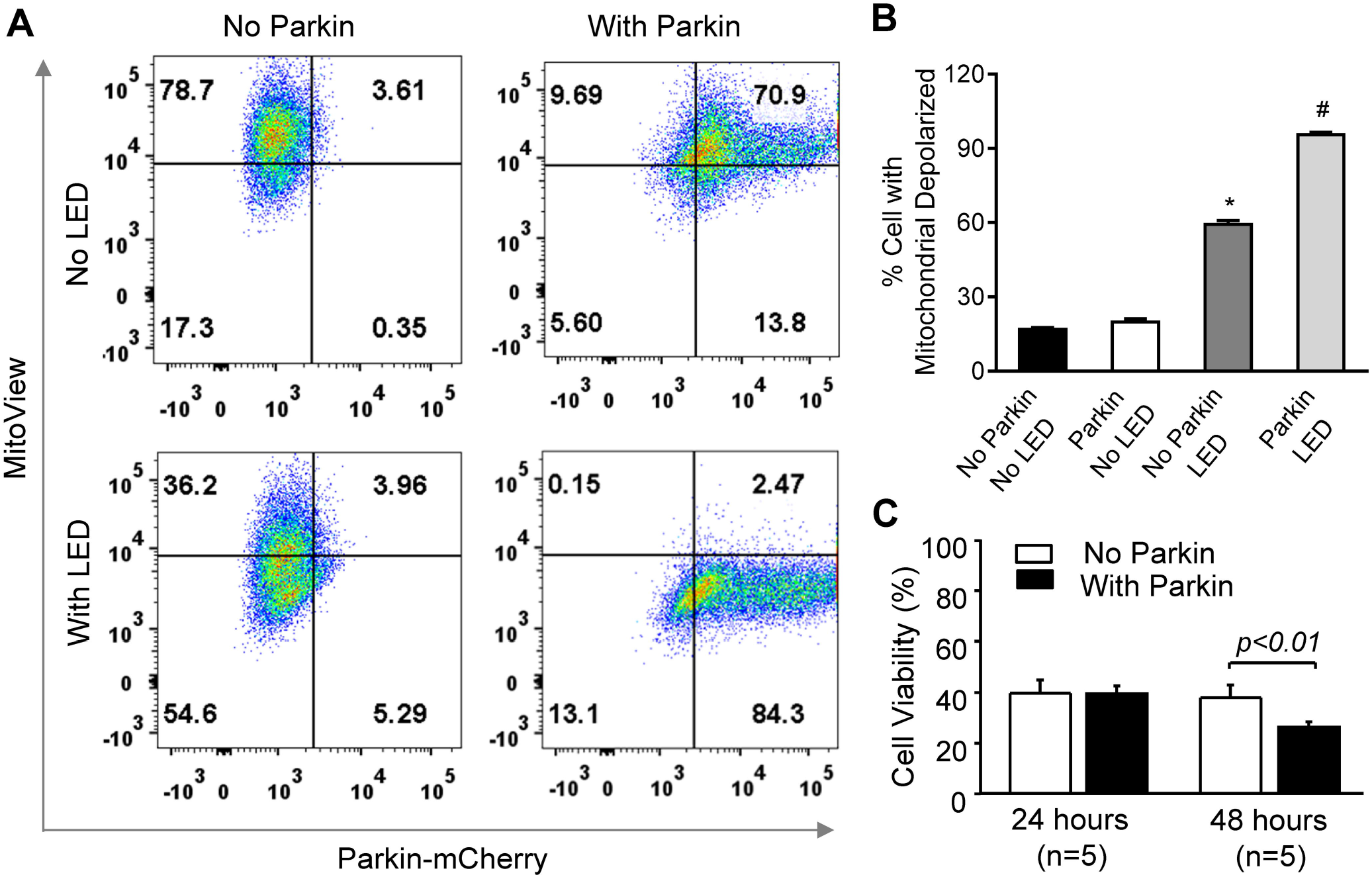
Optogenetic-mediated mitochondrial preconditioning protects cells against apoptotic cell death. A): Representative images of Hoechst and Propidium Iodide (PI) staining in cultures illuminated by LED light of moderate irradiance (4 mW/mm^2^ for 6 hours); B) Representative images of Hoechst and PI staining in cultures illuminated with mild (0.2 mW/mm^2^ for 2 hours) LED followed by moderate (4 mW/mm^2^ for 6 hours) LED illumination; C): Summary of cell viability in the unilluminated, illuminated only, and preconditioned groups; D-F): Representative confocal images showing MitoView fluorescence in the un-illuminated (D), illuminated only (E) and preconditioned and illuminated (F) cell cultures; and G): Normalized mitochondrial membrane potential (ΔΨ_m_, measured by MitoView) in different groups. *: P<0.05 *vs*. No illumination. #: P<0.05 *vs*. LED illumination. 3 cell cultures were examined for each group.

## DISCUSSION

Although optogenetics has been widely used to control the activity of excitable cells *in vitro* and *in vivo* (25, 44, 45), application of this innovative technique to manipulate intracellular organelles such as mitochondria has been limited until recently (46, 47). One challenge to developing a mitochondrial-based optogenetic approach is targeted expression of light-gated channel proteins in the highly folded IMM. In the present study, we have examined the capability of several commonly used MLS, including *mito*, to target the heterologous ChR2 protein to mitochondria. While Tkatch *et al.* have reported that tandem repeats of the *mito* sequence can target ChR2 to mitochondria (46), triplicate *mito* sequences led to little ChR2 mitochondrial localization in our hands. Instead, we provide evidence that the ABCB10 MLS can effectively translocate ChR2 to the IMM in a variety of cell types including H9C2, HeLa and cardiomyocytes. In addition, with the ABCB MLS the plasma membrane targeting sequence of ChR2 does not need be removed for mitochondrial localization. We further demonstrate that the expressed ChR2 proteins form functional cationic ion channels in the IMM, enabling targeted induction of mitochondrial depolarization by blue light. We speculate that although ChR2 channels are non-selective and purely passive and thus would allow the transport of any ions across the inner membrane upon activation following the electrochemical gradient, it is influx of protons, instead of influx of other cations or efflux of anions such as Cl^−^ and O_2_^.−^, that is responsible for the light-induced mitochondrial depolarization, based on the relative permeability of cations (H^+^ ≫ Na^+^ > K^+^ ≫ Ca^2+^ (48)), limited permeability to Cl^−^ (17), and lack of effect of an mPTP blocker (Figure 3). We confirm the optogenetic-mediated ΔΨ_m_ dissipation by measuring ΔΨ_m_ using confocal live cell imaging and flow cytometry, as well as by the translocation of Parkin from the cytosol to mitochondria, which is a well-established marker of mitochondrial depolarization.

The advantages of the optogenetic-based approach over the conventional pharmacological methods to induce mitochondrial depolarization include high spatial resolution *via* optics and specific targeting of ChR2 using cell type-specific promoters or vectors. The stimulus can be spatially controlled to a high extent, particularly at subcellular dimensions, due to its optical nature. Thus, optogenetic-based light illumination would allow controllable manipulation of individual mitochondria or groups of mitochondria within a single cell. The capability of inducing mitochondrial membrane permeability in specific mitochondria with light can enable novel research, such as investigating how ΔΨ_m_ impacts mitochondrial dynamics and network properties, Ca^2+^ handling, motility, mitophagy, or ROS-dependent signal propagation within or across cells. Spatial resolution is especially important for studies of neurons, which have much of their mitochondrial mass located in distal axons or dendrites. There has been considerable debate regarding how neurons handle damaged or dysfunctional mitochondria with some studies concluding that Parkin-mediated mitophagy only occurs near mature lysosomes in the cell body and that retrograde transport is required for degradation of distal mitochondria, while others concluding that Parkin-mediated mitophagy occurs in distal axons without retrograde transport to the soma (49, 50). Moreover, there may be distinct mechanisms by which neurons handle proximal vs. distal dysfunctional mitochondria and it is important to better define these mechanisms given that mitochondrial dysfunction is implicated in many neurodegenerative diseases and that degeneration of distal axon terminals may be one of the earliest stages and a better therapeutic target (51). The methods described here are ideally suited for investigating this because illumination can selectively depolarize only distal or only proximal mitochondria *in vitro* or *in vivo* using fiber optics.

In addition to high spatial resolution, this optogenetics approach provides specific targeting of tissues or cell types. Chemical uncoupling agents, such as the ionophores FCCP and CCCP, are widely used for the induction of mitochondrial depolarization. However, interpretation of the results can be confounded, especially in intact cells, because of their known off-target effects. For instance, it has been reported that both FCCP and CCCP inhibit the movement of mitochondria in neurites of chick sensory neurons in culture (52). Studies have also shown that FCCP mediates conductance of several ions (e.g., H^+^, Na^+^ and Cl^−^) in the plasma membrane (53), affecting membrane electrical characteristics (such as causing cell depolarization) (54),(55). On the contrary, the mitochondrial optogenetics approach shown here allows direct light-elicited ΔΨ_m_ depolarization by targeted expression of ChR2 proteins in the IMM. Moreover, compared with the conventional photoactivation method that requires high energy laser flashes, optogenetic-mediated ΔΨm depolarization requires very low energy (on the order of mW/mm^2^), as optogenetics is not simply photoexcitation of targeted cells; rather, it delivers gain of function of precise events. The low intensity light illumination would minimize side effects (such as photobleaching) on the illuminated cells or the proximal un-illuminated cells.

Apoptosis is a process of programmed cell death that occurs in multicellular organisms. While it is well established that mitochondria play key roles in activating programmed cell death, the underlying detailed mechanisms remain incompletely understood. For instance, the role of inner membrane permeabilization in the execution of pro-apoptotic protein (e.g. cytochrome C) release from the intermembrane space into the cytosol is a subject of ongoing controversy. Some studies show that mitochondrial depolarization and IMM permeabilization are required for cytochrome C release and apoptotic cell death (56). However, others suggest that cytochrome C release can occur in the absence of ΔΨ_m_ depolarization (57), mitochondrial depolarization may not be involved in the activation of apoptosis (58), or mPTP opening is a consequence of apoptosis (56). The disparity may be attributed to the methods used in different studies, such as pharmacological intervention versus genetic manipulation. In the present study, using the mitochondrial optogenetics approach we demonstrate that increased IMM permeability alone is sufficient to trigger apoptotic cell death, which is time and (light) dosage dependent. The activation of intrinsic apoptosis is confirmed by induction of cytochrome C release and prevention of cell death by caspase inhibition (Figure 5). Importantly, blocking mPTP did not prevent the light-induced cell death, which provides additional evidence that the expressed ChR2 formed functional cationic pores in IMM. Intriguingly, we showed that mild short-term light illumination ameliorates apoptotic cell death in ABCB-ChR2-expressing cells, supporting the notion of mitochondrial preconditioning (Figure 8). Thus, this innovative mitochondrial optogenetics technique can be used to effectively control IMM depolarization and differentially modulate cell fate, a capability not possessed by the existing pharmacological approaches.

Loss-of-function mutations in Parkin have been causally linked to early-onset Parkinson’s disease (59). In addition, studies have shown that Parkin overexpression can prevent cell death in response to a variety of stimuli (60–62). Thus, considerable enthusiasm has emerged recently regarding the potential of Parkin as a therapeutic target. However, here we surprisingly observe that activation of Parkin exacerbates light-induced mitochondrial depolarization and cell death. While our findings appear contradictory to the conventional notion that Parkin is cytoprotective, the pro-apoptotic effect of Parkin has been reported in two recent studies (42, 43). The paradoxical role of Parkin appears attributed to the nature of the stressor to induce cell death, i.e., whether the stimulus induces mitochondrial depolarization and activation of Parkin. For instance, Carroll *et al.* (42) have shown that Parkin overexpression enhances HeLa cell apoptosis induced by mitochondrial depolarizing agents such as CCCP and valinomycin but has no effects on cell death induced by other stimuli that fail to activate Parkin. Similarly, Zhang *et al.* reported that Parkin activation significantly exacerbates valinomycin-induced apoptosis but does not promote CCCP-induced cell death in HeLa cells (43). They further demonstrated that the level of Parkin recruitment-induced Mcl-1 degradation and polyubiquitination is a key factor determining whether Parkin-dependent mitophagy versus apoptotic cell death is activated. While the reason underlying the discriminate effect of Parkin on the fate of CCCP-treated cells in those two studies requires further investigation, the severity of mitochondrial damage might be a critical factor. If mitochondrial damage is too severe to be repaired by mitophagy, the PINK1/Parkin pathway may promote apoptosis. The effect of Parkin on cell death decisions may also depend on the level of its activation: with transient activation inducing mitophagy to repair mitochondrial networks and sustained activation triggering apoptosis to eliminate damaged cells. Our findings warrant consideration when designing Parkin-based therapeutic strategies.

One limitation of the developed mitochondrial optogenetics technique is that although it enables targeted mitochondrial depolarization at high spatial (cellular or subcellular) resolution, the time resolution of light-induced mitochondrial depolarization is much less. Similar to findings shown in a recent study (46), optogenetic-induced mitochondrial depolarization takes several minutes to accomplish. Using other rhodopsin protein variants with higher conductance or greater light sensitivity may help to achieve better temporal control of mitochondrial depolarization (63). The lack of mitochondrial repolarization may be partially due to the slow deactivation of ChR2 channels. Tkatch *et al.* (46) showed that optogenetic-induced mitochondrial depolarization can recover slowly in cells expressing the ChR(SSFO) mutant. Alternatively, co-expressing Channelrhodopsin and Bacteriorhodopsin or Archaerhodopsins in the IMM and stimulating cells with alternating blue and yellow light might achieve reversible control of ΔΨ_m_. The energetic states or types of cell may also affect the feasibility of inducing faster or more reversible mitochondrial depolarization. For instance, mitochondrial oscillations can be readily triggered in isolated guinea pig cardiomyocytes (7, 9), but it is much more difficult in isolated mouse cardiomyocytes.

## CONCLUSIONS

We have developed and characterized a mitochondrial optogenetics approach to induce targeted mitochondrial depolarization, which provides an alternative to the traditional optical or pharmacological means of modulating MMP and ΔΨ_m_ to understand their roles in determining cell fate. We expect the research tools shown here and the knowledge acquired will facilitate the design of novel optogenetic-based therapeutic strategies for the treatment of diseases involving mitochondrial dysfunction.

## ACKNOWLEDGEMENTS

The authors thank Drs. Kah Yong Goh, Jiajia Song and Qince Li for their technical support in cell death assay and immunostaining. This work was supported by National Institute of Health (NIH) R21 HL127599 and R01 HL121206 (to L. Z.) and American Heart Association (AHA) Predoctoral Fellowship 18PRE34060188 (to P. E.).

