## Supplemental figures for "Precisely control mitochondria with light to manipulate cell fate decision"

**Running title**: Optogenetic control of mitochondria

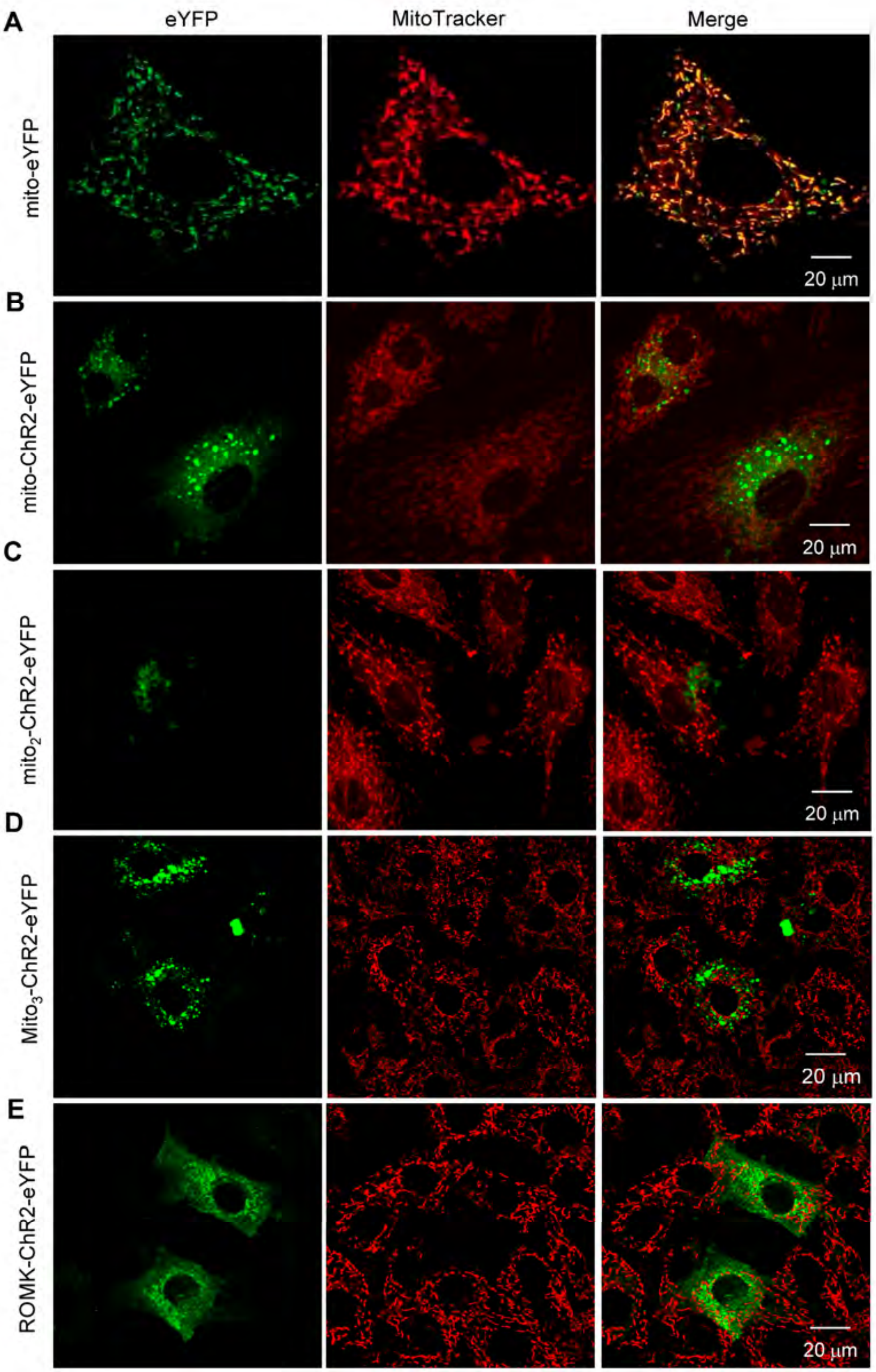


**Figure S1**. Representative confocal images showing intracellular localization of mito-eYFP (A), mito-ChR2-eYFP (B), mito$_2$-ChR2-eYFP (C), mito$_3$-ChR2-eYFP (D), and ROMK-ChR2-eYFP (E) in H9C2 cells. Mitochondria were stained with fluorescent dye MitoTracker Deep Red (250 nM). Yellow indicates overlay of eYFP and the mitochondrial marker.

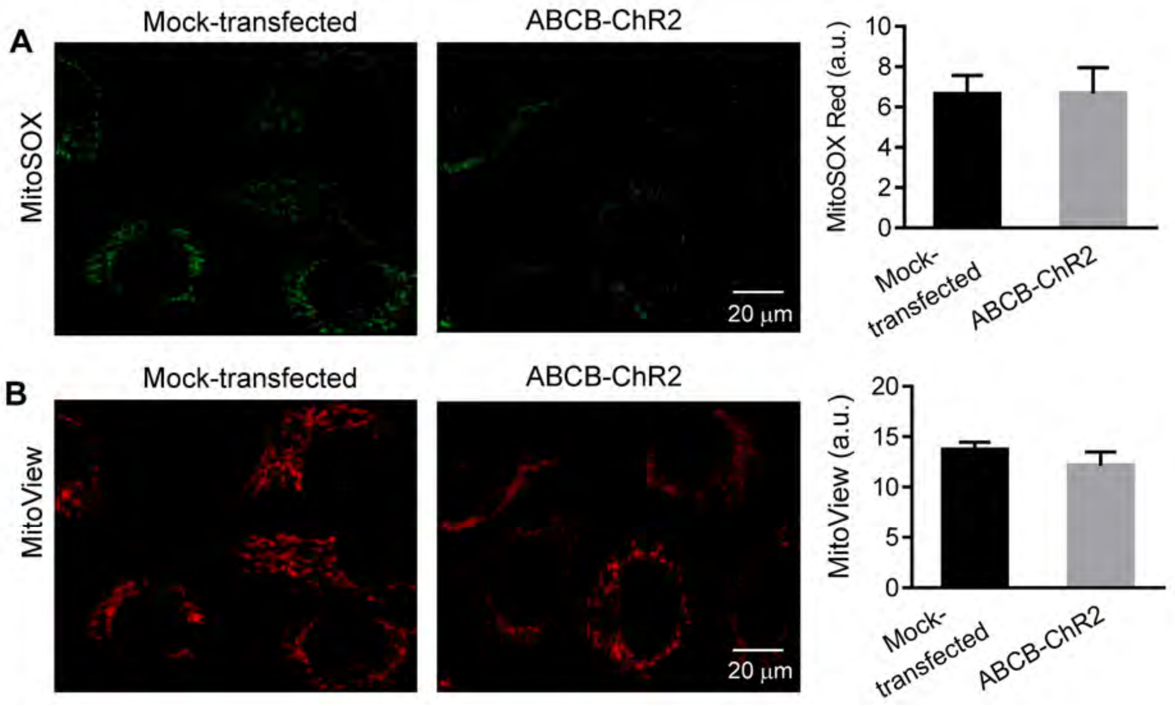


**Figure S2.** Repesentative confocal images showing mock-transfected and ABCB-ChR2-transfected HeLa cells stained with MitoSox (1 μM) (A) and MitoView (25 nM) (B). Data analysis revealed that transfection and expression of ABCB-ChR2 had little effect on mitochondrial ROS and membrane potential in HeLa cells. n=4 each group.

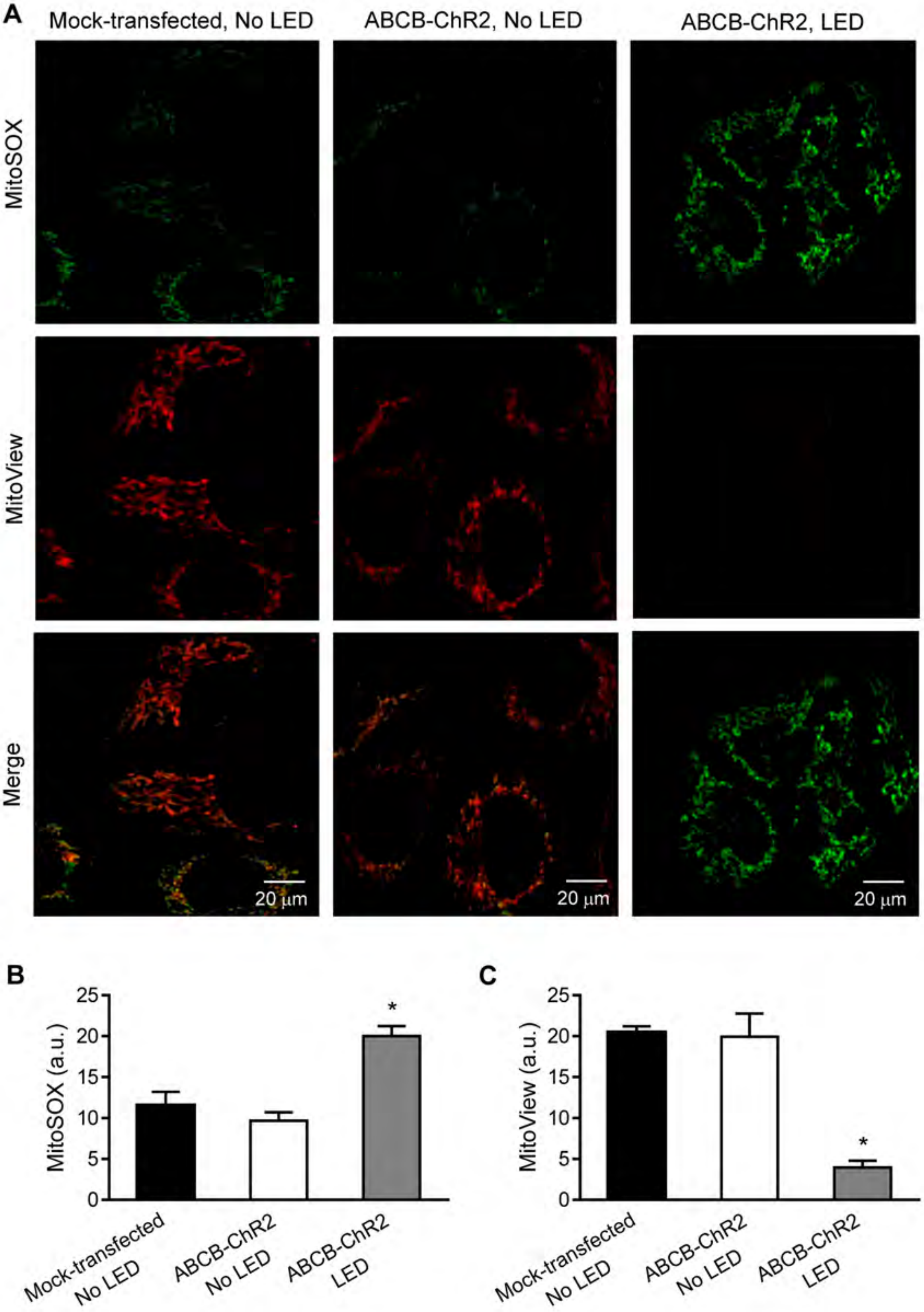


**Figure S3.** A): Representative confocal images of HeLa cells stained with MitoSox (green) and MitoView (red) showing that blue LED pulse illumination caused increased ROS production and loss of mitochondrial membrane potential in ABCB-ChR2-expressing cells; B): Analytical data of MitoSOX; and C): Analytical data of MitoView.

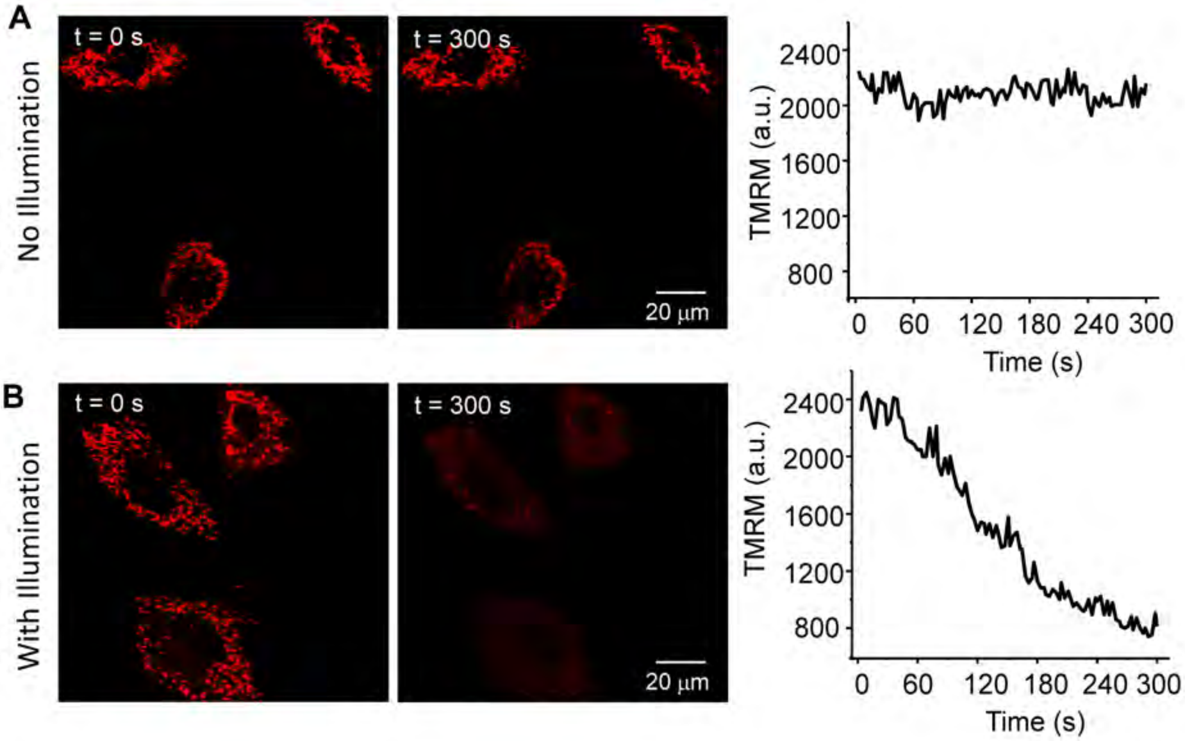


**Figure S4.** Confocal recording of mitochondrial membrane potential in ABCB-ChR2-eYFP-expressing HeLa cells. Cells were stained with TMRM (20 nM) and then imaged for 300s in the absence (A) or presence (B) of 475nm LED (5 mW/mm$^2$) pulse illumination. Change of TMRM fluorescence was analyzed offline using imageJ. Three experiments were repeated.

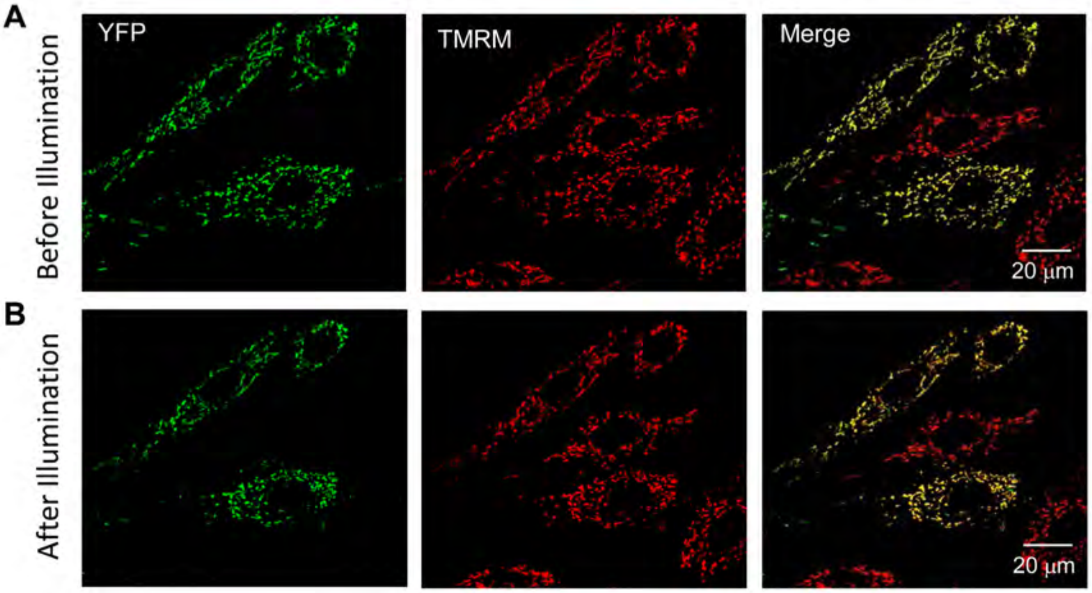


**Figure S5.** Confocal images of mitochondrial membrane potential in ABCB-eYFP-expressing HeLa cells before and after light illumination. Cells were stained with TMRM (20 nM) and then imaged for 300s in the presence of 475nm LED (5 $mW/mm^2$) pulse illumination. Top: eYFP and TMRM before illumination, lower: eYFP and TMRM after illumination. Three experiments were repeated.

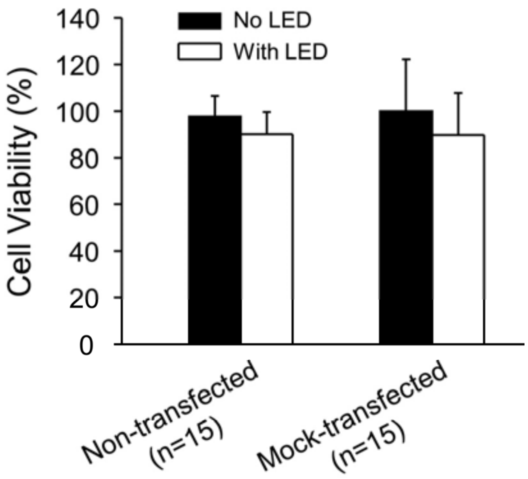


**Figure S6:** The LED illumination same as that used in figure 3 did not elicit cell death in the non-transfected and mock-transfected HeLa cells. n represents the number of cell cultures.

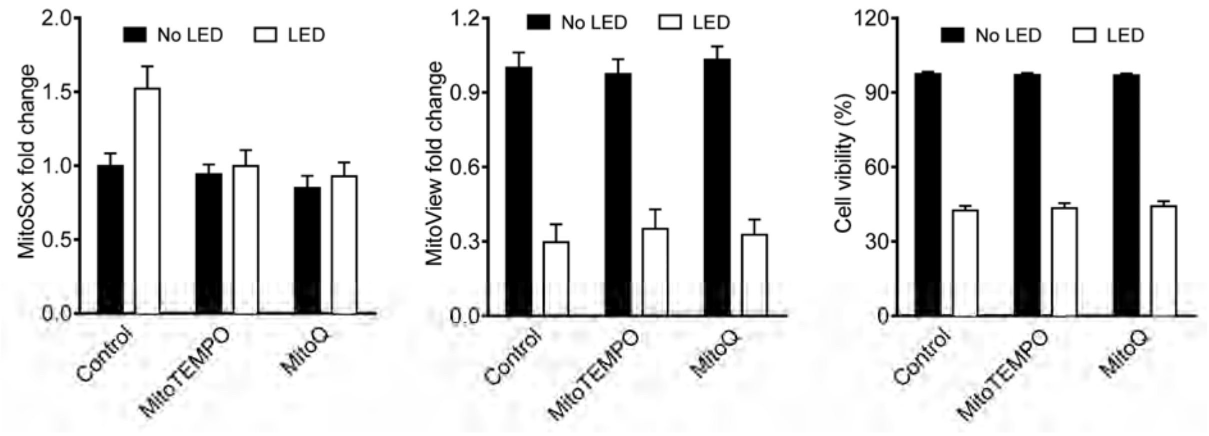


**Figure S7.** Mitochondrial-targeted antioxidant MitoTEMPO and MitoQ reduced ROS accumulation (left) but had little effect on mitochondrial membrane potential (middle) and cell viability (right) in ABCB-ChR2-expressing cells exposed to light illumination (0.5 $mW/mm^2$, 24 hours). ROS was measured with MitoSox fluorescent dye, mitochondrial membrane potential was measured with MitoView fluorescent dye, and cell viability was measured by trypan blue method. n=4 each group.

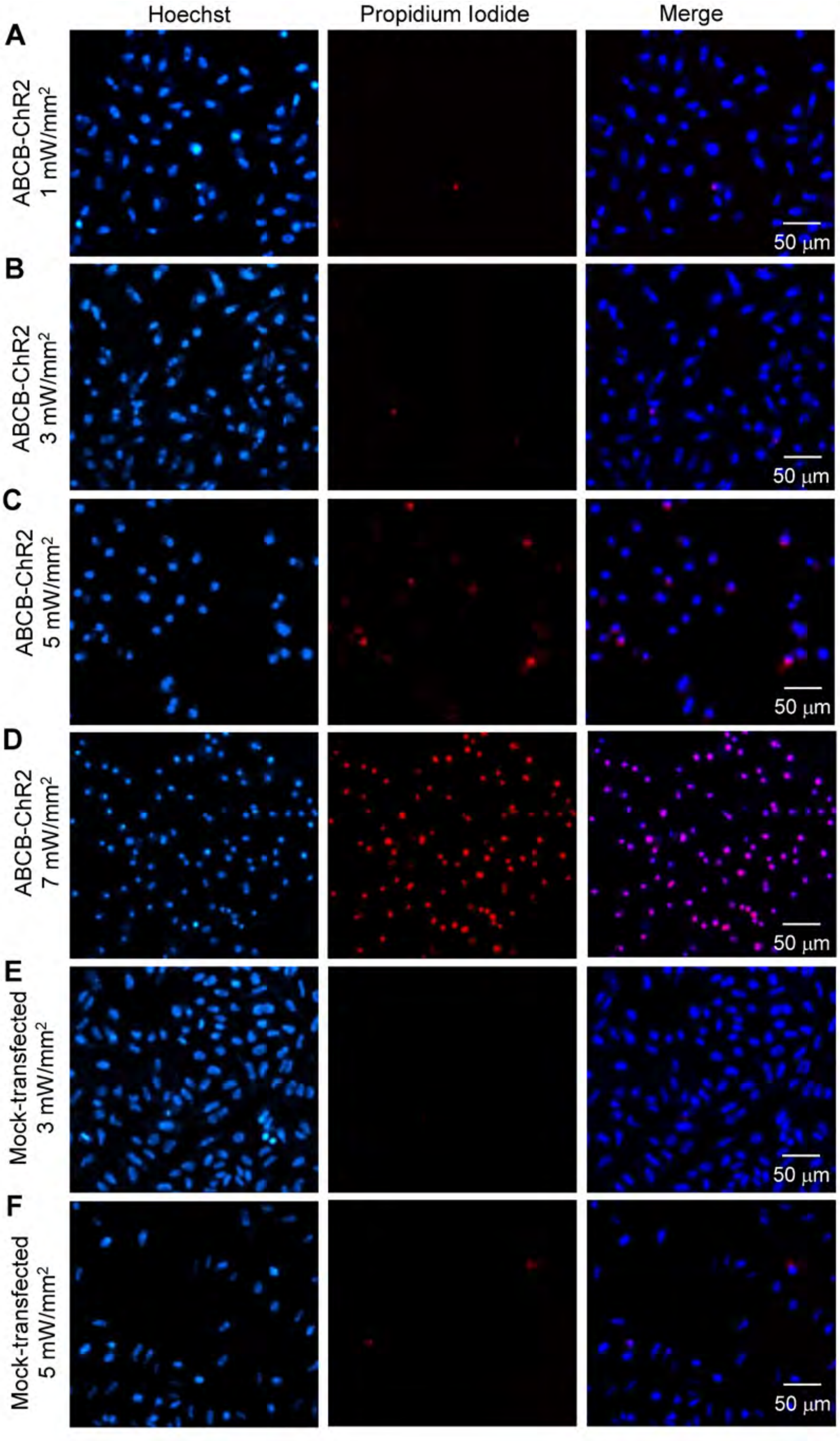
Hoechst
Propidium Iodide
Merge
A
ABCB-ChR2
1 mW/mm²
50 μm
B
ABCB-ChR2
3 mW/mm²
50 μm
C
ABCB-ChR2
5 mW/mm²
50 μm
D
ABCB-ChR2
7 mW/mm²
50 μm
E
Mock-transfected
3 mW/mm²
50 μm
F
Mock-transfected
5 mW/mm²
50 μm

**Figure S8.** Light irradiance dependence of optogenetics-mediated cell death. After 6 hours of LED illumination at various intensities, ABCB-ChR2-expressing (A-D) or mock-transfected (E and F) cells were double-stained with Hoechst (blue) and propidium iodide (red) and subjected to fluorescent microscopy imaging.

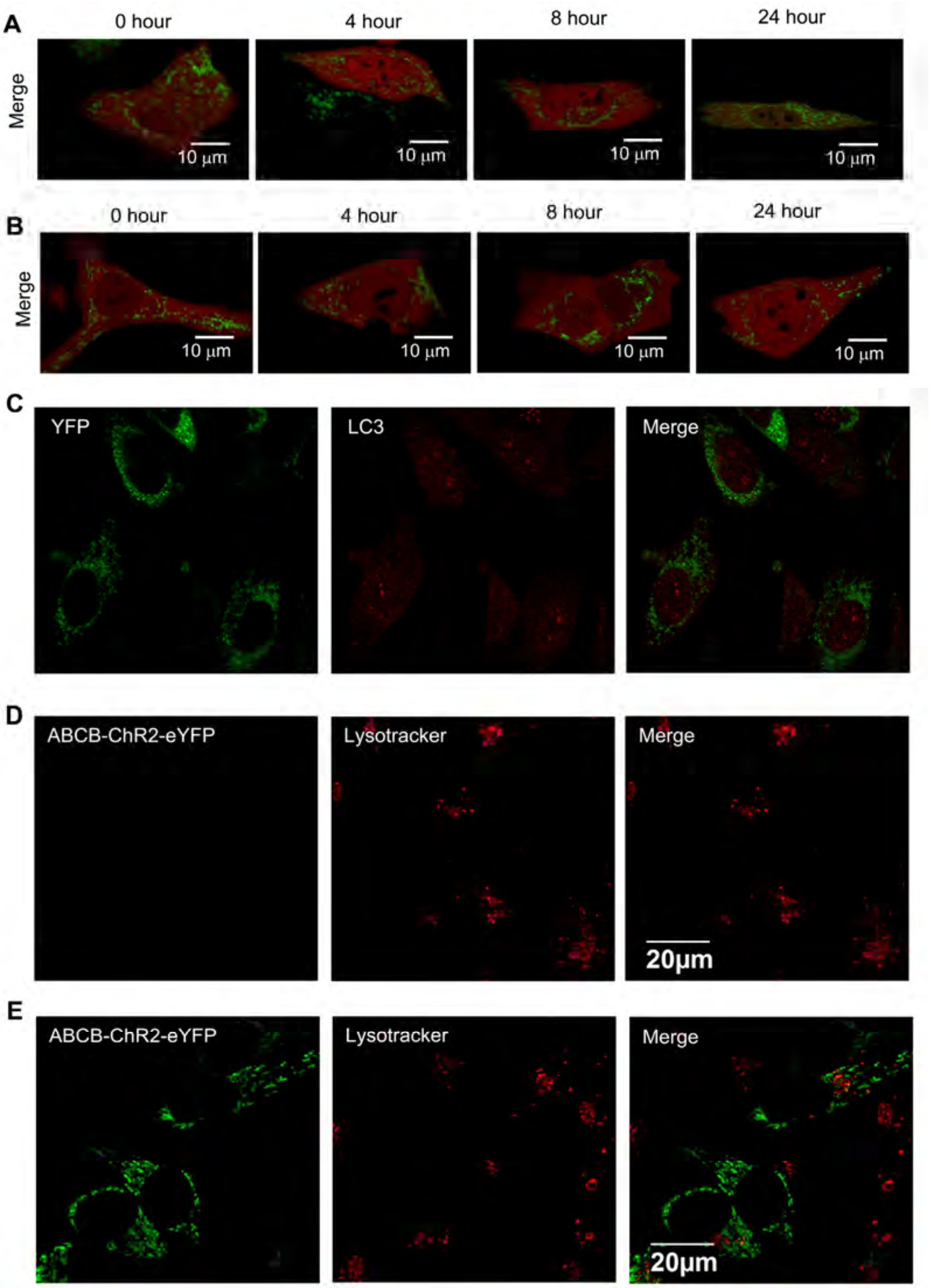


**Figure S9.** A and B: Representative confocal imagines of light illuminated cells co-expressing ABCB-eYFP and Parkin-mCherry (A) and un-illuminated cells co-expressing ABCB-ChR2-eYFP and Parkin-mCherry (B). Images were taken at 0, 4, 8, and 24 hours during light illumination. Red represents Parkin-mCherry, green represents eYFP, and merge represents Parkin accumulation in mitochondria. C: Confocal image showing lack of LC3 aggregation in the un-illuminated ABCB-ChR2-eYFP expressing cells. D and E: Confocal images showing lack of YFP and lysosome colocalization in the mock-transfected cells (D) and un-illuminated ABCB-ChR2-eYFP expressing cells (E).

**Video S1 for Figure 3.** Blue LED illumination caused targeted (zone 1 in figure 2) mitochondrial depolarization in H9C2 cells expressing ABCB-ChR2-eYFP. TMRM fluorescence remained unchanged in zone 2 cells that were not illuminated by blue light.

**Video S2 for Figure S4.** A. In the absence of blue light illumination, the imaging laser (543 for TMRM) did not affect the fluorescence of TMRM in H9C2 cells expressing ABCB-ChR2-eYFP. B: Concurrent blue LED illumination caused mitochondrial depolarization in H9C2 cells expressing ABCB-ChR2-eYFP.

**Video S3 for figure S5.** In the absence of ChR2 expression, LED illumination did not affect TMRM fluorescent intensity in H9C2 cells expressing ABCB-eYFP.
